# Factors associated with opioid cessation: a machine learning approach

**DOI:** 10.1101/734889

**Authors:** Jiayi W. Cox, Richard M. Sherva, Kathryn L. Lunetta, Richard Saitz, Mark Kon, Henry R. Kranzler, Joel E. Gelernter, Lindsay A. Farrer

**Affiliations:** Section of Biomedical Genetics, Boston University School of Medicine, Boston, MA, USA; Department of Biostatistics, Boston University School of Public Health, Boston, MA, USA; Department of Community Health Sciences, Boston University School of Public Health, Boston, MA, USA; Department of Mathematics and Statistics, Boston University College of Arts & Sciences, Boston, MA, USA; Department of Psychiatry, Perelman School of Medicine, University of Pennsylvania, Philadelphia, PA, USA; Department of Psychiatry, Yale School of Medicine, West Haven, CT, USA; Department of Genetics and Genomics, Boston University School of Medicine, Boston, MA, USA

**Keywords:** opioid use disorder, opioid cessation, opioid abstinence, machine learning, AI, diagnostic questionnaire, feature selection, outcome prediction

## Abstract

**Background and Aims:** People with opioid use disorder (OUD) can stop using opioids on their own, with help from groups and with treatment, but there is limited research on the factors that influence opioid cessation.

**Methods:** We employed multiple machine learning prediction algorithms (LASSO, random forest, deep neural network, and support vector machine) to assess factors associated with ceasing opioid use in a sample comprised of African Americans (AAs) and European Americans (EAs) who met DSM-5 criteria for mild to severe OUD. Values for several thousand demographic, alcohol and other drug use, general health, and behavioral variables, as well as diagnoses for other psychiatric disorders, were obtained for each participant from a detailed semi-structured interview.

**Results:** Support vector machine models performed marginally better on average than those derived using other machine learning methods with maximum prediction accuracies of 75.4% in AAs and 79.4% in EAs. Subsequent stepwise regression analyses that considered the 83 most highly ranked variables across all methods and models identified less recent cocaine use (p<5×10^−8^), a shorter duration of opioid use (p<5×10^−6^), and older age (p<5×10^−9^) as the strongest independent predictors of opioid cessation. Factors related to drug use comprised about half of the significant independent predictors, with other predictors related to non-drug use behaviors, psychiatric disorders, overall health, and demographics.

**Conclusions:** These proof-of-concept findings provide information that can help develop strategies for improving OUD management and the methods we applied provide a framework for personalizing OUD treatment.

## Introduction

Misuse of prescription opioids and illicit opioids are a significant global problem that affects the health and economic welfare of individuals, families, and society. The opioid overdose rate has quadrupled since 1991(1). In 2017, more than 47,000 Americans died of an opioid overdose, and 36% of these deaths involved prescription opioids(2). A major goal of addressing opioid use disorder (OUD) is to achieve abstinence, or cessation, from illicit use and misuse. There is not a single, clinically accepted definition of cessation with regard to length of time without illicit use or prescribed opioids misuse. (3, 4). Previous studies identified age, employment status, and age at first drug use to be associated with successful completion of a treatment program (3). Other factors are likely to influence successful cessation. Based on studies of OUD severity (5–7), pain experiences, general health, use of antidepressants may also influence cessation. Delineation of these factors could inform efforts to help patients or be useful for patients to practice self-care. However, studies thus far have tended to focus on a small number of clinically relevant factors such as medication dose, duration and formulation (8–10). There are currently much larger epidemiological studies of OUD (11–13) with data on thousands of potential predictors that would enable a systemic hypothesis-free query to identify factors predicting opioid cessation.

Many statistical methods are limited in their ability to sort through large numbers of predictors(14). Data mining using machine learning, which is particularly well suited for identifying predictive factors among thousands of variables (15, 16), has successfully identified predictor variables for a diverse set of outcomes(17–21). Here, we applied multiple machine learning techniques to evaluate a large set of clinical, demographic, general health, and behavioral variables in a large, racially mixed cohort who were recruited for genetic studies of substance dependence and not necessarily treated for OUD to identify factors that influence opioid cessation (defined as self-reported last illicit opioid use and/or prescription opioids misuse > 1 year before the interview date). Identifying additional factors associated with cessation could support a personalized approach to improve the outcome of cessation attempts.

## Material and Methods

### Participants and Assessments

Participants for this study were selected from a cohort of 6,188 African Americans (AAs) and 6,835 persons of European Americans (EAs) who were recruited for genetic studies of opioid, cocaine, or alcohol dependence between 2000 and 2017 through advertisements and treatment clinics at the University of Connecticut Health Center, Yale University School of Medicine, the Medical University of South Carolina, the University of Pennsylvania, and the McLean Hospital(22, 23). This cohort included affected sibling pairs and additional family members, as well as unrelated cases and controls. Persons with schizophrenia or schizoaffective disorder were excluded(22, 23). Information about use of various substances, demographics, general health, behavior, and other psychiatric illnesses was obtained by interview using the Semi-Structured Assessment for Drug Dependence and Alcoholism (SSADDA) (11, 24). Substance use disorder (SUD) and psychiatric disorders were established according to DSM-IV criteria. Institutional review boards from each recruitment site and Boston University approved this study, and written informed consent was obtained from all participants.

### Opioid Cessation Definition

Participants who were eligible for this analysis met at least two DSM-5 criteria for OUD, corresponding to a lifetime diagnosis of mild to severe OUD. Current opioid cessation was determined by the response to the question, “When was the last time you used an opioid drug (including illicit methadone).” This question was asked as part of a series of items asked about illicit or non-prescribed use of opioids. Individuals who last used an opioid >1 year before the date of interview were considered to have achieved cessation and those whose last use of an opioid was <6 months before the interview date were classified as non-cessation. Persons who used opioids between 6 months and 1 year before the interview date were excluded from further analysis. Filtering steps yielding a sample of 1,192 AAs and 2,557 EAs for analysis is shown in Figure S1.

### Phenotype Data Processing

Preprocessing of 3,956 SSADDA variables was performed prior to machine learning analyses. Variables with narrative or invariable responses, containing redundant information (e.g., specific date of different episodes, drug names), and with a response rate <90% were removed. Missing values for binary and categorical variables were recoded as indicator variables to accommodate missing responses. Missing values for continuous variables were imputed to the population group mean value. Missing values for ordinal variables related to recency of drug use were coded as dummy variables to account for current of never use. Z-score normalization (mean of 0 and variance of 1) was applied to continuous variables within each population to minimize scaling issues. The number of variables remaining after these steps was 3,315 in AAs and 3,738 in EAs.

### Machine Learning Analyses

AAs and EAs were analyzed separately because they had different SSADDA completion rates and combining the two groups might lead to response bias. Variables were grouped into three nested sets defining three analytical models. Model 1 contained all variables except those related to recency of opioid use which are strongly correlated with cessation, thus with variable totals of 3,093 in AAs and 3,503 in EAs. Model 2 further excluded all opioid-related questions, leaving 2,863 variables in AAs and 3,252 in EAs. Model 3 further excluded all drug use variables, leaving only demographic, non-SUD diagnoses, behavioral, and general health variables (1,656 in AAs and 1,907 in EAs). Models were evaluated using four machine learning methods described in Supplemental Materials to identify variables predicting opioid cessation. We modeled different types of intra-variable relationships between predictors and the outcome using linear (LASSO(25) and linear support vector machine with recursive feature elimination [SVM](26)) and non-linear (random forest with recursive feature elimination [RF](27) and deep neural network with feature selection [DNN] (28)) techniques. These four methods were applied to capture predictive variables under different model assumptions and allow for different outcome-predictor relationships. Variables from each model that were associated with the highest accuracy reflected by either F1 score or area under the curve (AUC) and generated by each machine learning method were retained. The F1 score is a harmonic measure of precision (true positive / [true positive + false negative]) and recall (true positive / [true positive + true negative]),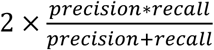 at a given case/control split, and AUC is an overall evaluation of model performance that accounts for the true positive and false positive rates for all possible diagnostic splits(29, 30). Both measurements were considered because of their popularity in the clinical setting(31). The F1 score was used to assess accuracy due to limitations of the AUC, which includes bias when performed on imbalanced datasets as well as impractical and uninterpretable split points for evaluation(29, 32).

### Statistical Methods for Testing the Association of Opioid Cessation with Phenotypic Variables

To determine which variables selected by the machine learning methods independently predict cessation, we applied different cutoffs for the importance measurement of each machine learning method, namely the odds ratio (OR) for LASSO, coefficient(33) denoted by weight for SVM, feature importance(33) for RF, and activation potential(28) for DNN. For LASSO, we chose variables that yielded ORs >1.05 or <0.95. We applied the following criteria for selecting variables from SVM and RF analyses depending on the number of variables (n) selected for each model: if n >200, the top 30% of variables measured by absolute weight in SVM or importance in RF were designated as high impact; if 100 < n < 200, the top 50% were selected; if n<100, all variables were designated as high impact. For DNN, all selected variables were designated “high impact”. Joint association tests were performed using bi-directional stepwise logistic regression that included 83 “high-impact” variables that were culled from three models across four machine learning methods in the AA and EA datasets. Variables that yielded the highest Akaike information criterion (AIC) with p <0.05 from bi-directional stepwise logistic regression were grouped into “drug related”, “behavioral”, “general health”, and “demographic” categories.

## Results

Characteristics of the study samples are shown in Table 1. The sample included 1,069 unrelated AAs and 2,252 unrelated EAs, as well as 123 AAs and 305 EAs participants who were members of families containing a pair of siblings both with either opioid or cocaine dependence. There is a higher proportion of females among individuals who ceased opioids in both AAs (OR=1.35, P=6.7 ×10^−3^) and EAs (OR=1.31, P=1.1 ×10^−3^) compared to those who did not cease. Furthermore, participants who ceased opioid use were older by an average of 3.18 years in the AA group (P=1.0 ×10^−10^) and 6.1 years in the EA group (P=2.2 ×10^−16^) than those who did not cease. The mean number of lifetime DSM-5 OUD criteria met among those who ceased opioid use was not significantly different from those who did not cease.

**Table 1.** Participant characteristics.

|  |  | Time since last use |  |
| --- | --- | --- | --- |
|  |  | ≤ 6 month (not cease) | >1 year (ceased) |
| AAs<br>(N=1192) | Total N (% female) | 701 (33.5%) | 491 (40.5%) |
| | Age (Mean ± SD) | 42.6 ± 8.5 | 45.6 ± 8.3 |
| | OUD Symptom Counts (Mean ± SD) | 7.8 ± 2.4 | 7.6 ± 2.5 |
|  | # of families (N in families) | 35 (76) | 23 (47) |
| EAs<br>(N=2557) | Total N (% female) | 1714 (34.4%) | 843 (40.6%) |
| | Age (Mean ± SD) | 34.4 ± 10 | 40.5 ± 10.3 |
| | OUD symptom Counts (Mean ± SD) | 8.8 ± 1.9 | 8.4 ± 2.3 |
|  | # of families (N in families) | 114 (241) | 31 (64) |

AAs: African Americans, EAs: European Americans, OUD: opioid use disorder

### Feature selection

The F1 score was generally higher across models in both AAs and EAs using SVM than the other machine learning algorithms (Figure S2), although the differences in F1 score across methods were generally small, especially for models 1 and 2. A detailed discussion of the performance of each method for the three models is provided in Supplemental Materials and Figure S2.

Figure S3 shows the overlap of high impact variables chosen by the four machine learning methods. LASSO “high impact” variables almost entirely overlap with those from the other methods, while DNN-selected variables overlap the least with other method-selected variables. The majority of variables selected by non-LASSO methods are unique to those methods, however, there was high overlap in “high impact” variables selected by SVM and RF. Age was among the five top-ranked variables consistently identified by each method for each model in both AAs and EAs (Table S1).

### Factors Associated with Opioid Cessation

Stepwise regression analysis that considered independent predictors among 83 “high impact” variables culled from all models and machine learning methods (Table 2) revealed that age was one of the most significant predictors of opioid cessation in both AAs (OR=2.44 per standard deviation (SD) change, P=1.41×10^−12^) and EAs (OR=2.00 per SD change, P=5.74×10^−9^). Variables related to drug use comprised over 50% of the nominally significant predictors of opioid cessation in AAs (29 of 41) and EAs (27 of 50). Recency of last cocaine injection in AAs (OR=2.30 per level change, P = 9.11×10^−6^) or use in EAs (OR=1.91 per level change, P=3.30×10^−15^) were also among the most significant predictors. The number of years using heroin in AAs (OR=0.55 per SD change, P=5.78×10^−6^) and age at first heavy opioid use in EAs (OR=0.56 per SD change, P=2.67×10^−12^) were also significant predictors. Greater odds of cessation in AAs were associated with variables related to use of other drugs including “less recent of hving last alcohol symptoms” (OR=1.45 per level change, P=2.84×10^−3^ and “had 2 marijuana symptoms lasting >1 month” in AAs (OR=2.13, P=4.83×10^−3^). Drug use-related variables associated with cessation in EAs included less recent of last having alcohol symptoms (OR=1.34 per level change, P=4.13×10^−5^), “able to cut down smoking” (OR=1.28, P=3.69×10^−2^), and “marijuana interfered with work or home activities” (OR_EA_=1.67, P_EA_=1.61×10^−3^). The Fagerstrom Test for Nicotine Dependence (FTND) item “smoked more frequently after waking up” (OR=0.57, P=7.76×10^−3^) and having attended a self-help group for OUD (OR=0.57, P=1.41×10^−2^) were associated with decreased odds of cessation in AAs, whereas in EAs, starting attendance at an OUD self-help group sooner (OR=1.28 per level change, P=2.4×10^−3^) and recent attendence at a cocaine self-help group meeting (OR=0.76 per level change, P=3.27×10^−4^) increased the odds of cessation.

**Table 2.**
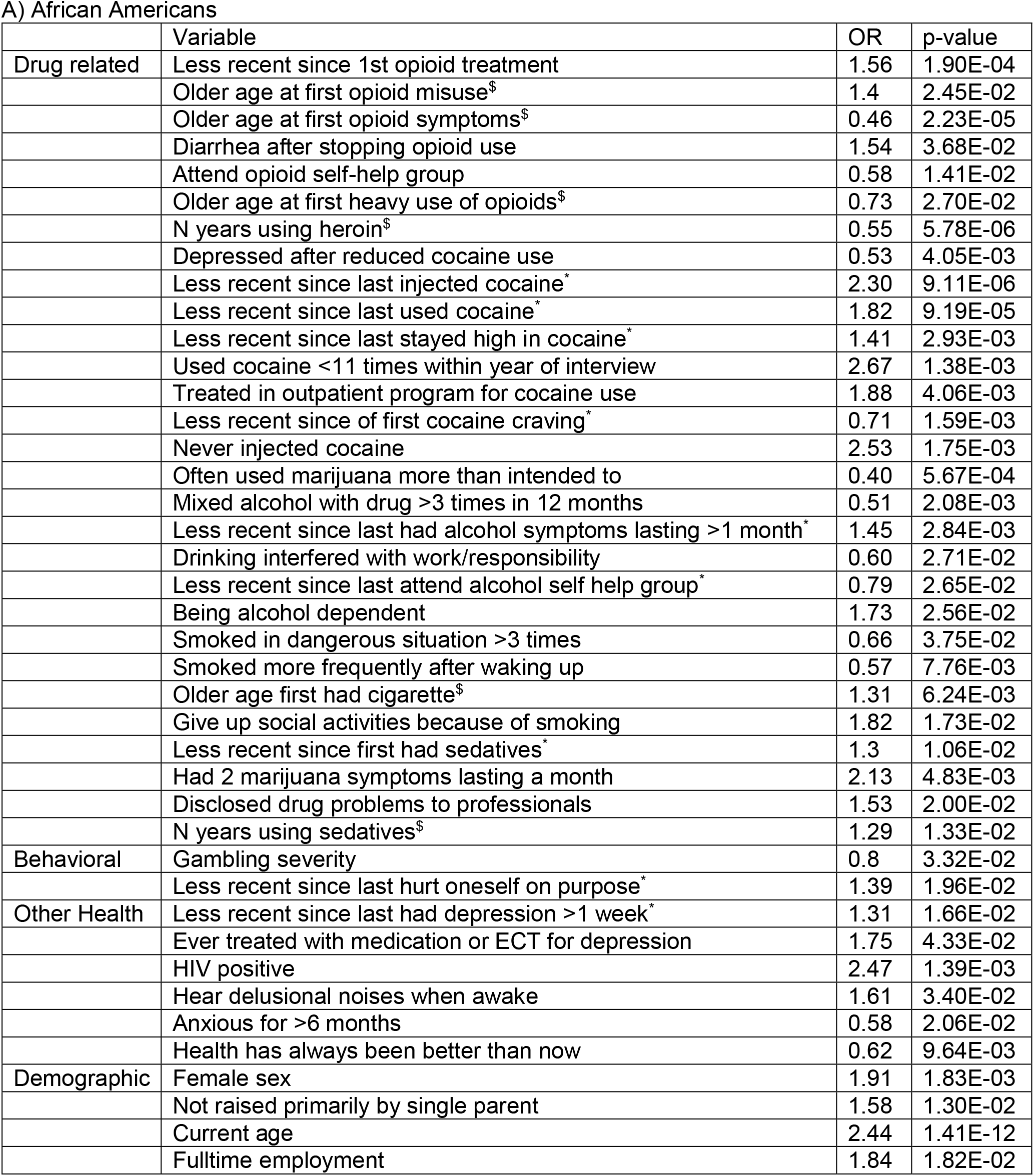
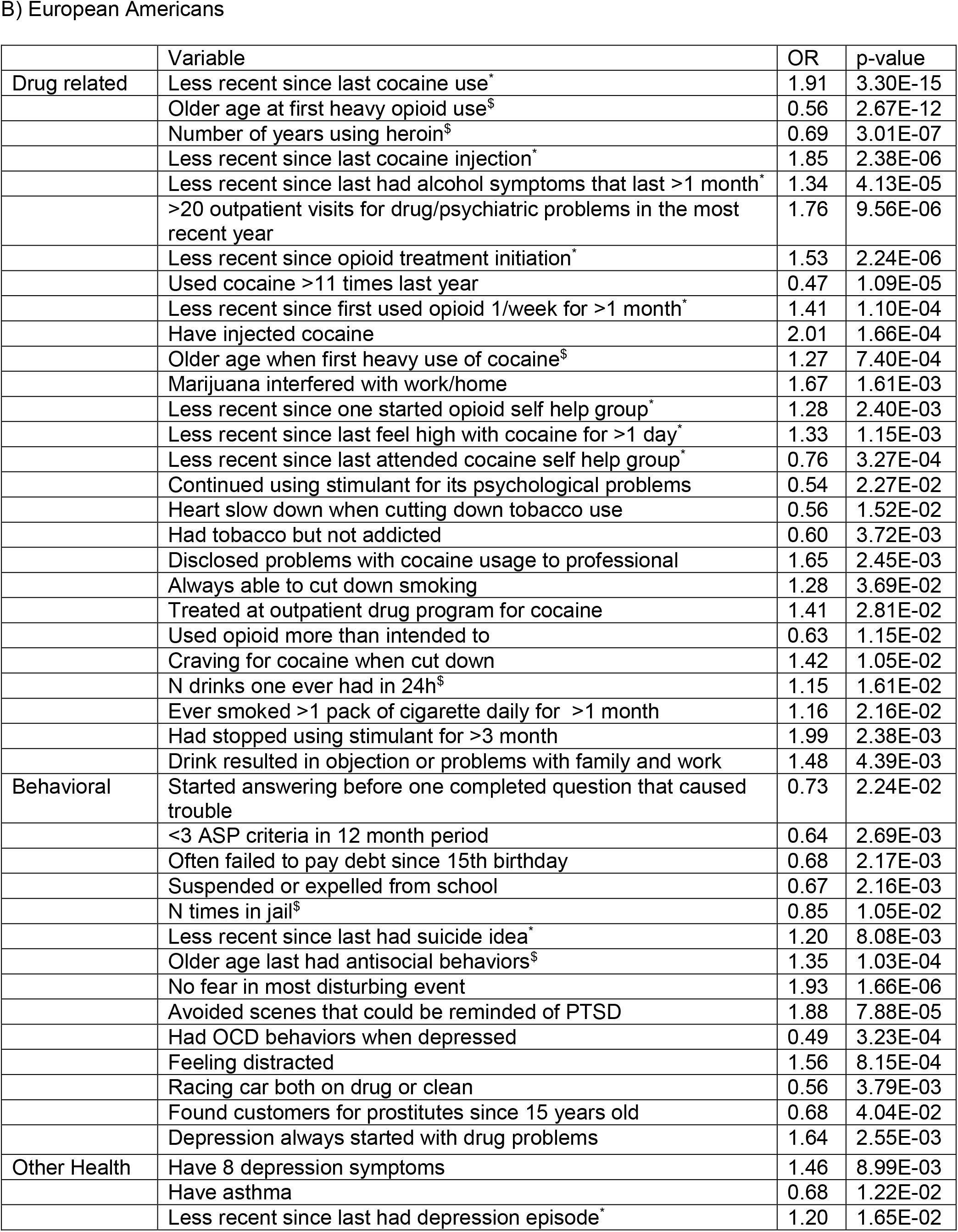
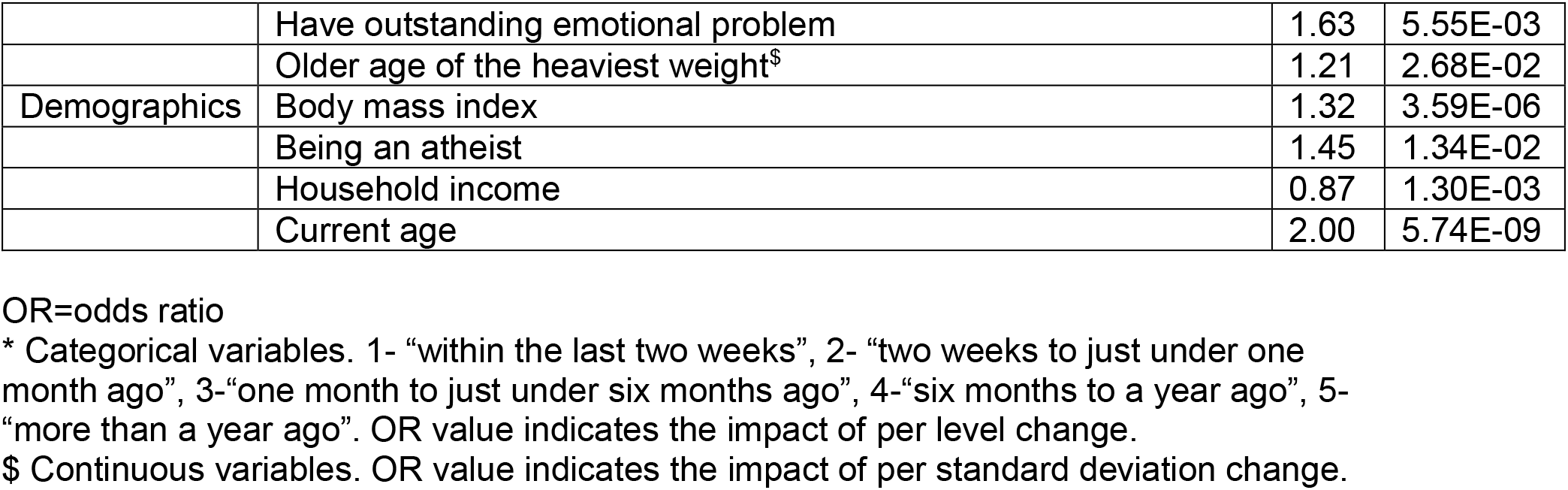
Stepwise regression results using all significant variables from all machine learning methods and models in (A) African Americans and (B) European Americans.

Among behavior-related variables, suicidal ideation or self harm was significantly associated with higher odds of cessation in both AAs (OR=1.39, P=1.96×10^−2^) and EAs (OR=1.2, P=8.08×10^−3^). In EAs, recovering from an event causing PTSD asssed by the question “No fear in most disturbing event” (OR=1.93, P=1.66×10^−6^) and no recent antisocial behavior episodes (OR=1.35 per SD change in age, P=1.03×10^−4^) were also associated with increased odds of cessation.

Several other health-related variables including past depressive episode lasting >1 week in AAs (OR=1.31, P=1.66×10^−2^), having drug-use associated depression in EAs (OR=1.64, P=2.55×10^−3^), body mass index in EAs (OR=1.32 per SD change, P=2.58×10^−6^), and being HIV positive in AAs (OR=2.47, P=1.39×10^−3^) were significantly predictive of cessation, whereas asthma (OR=0.68, P=1.22×10^−2^) significantly lowered the odds of cessation in EAs.

Other significant predictors of opioid cessation that were identified by this analysis include being employed (OR=1.84, P=1.82×10^−2^) and not having been raised by a single parent (OR=1.58, P=1.3×10^−2^) in AAs, and lower household gross income (OR=0.87 per level change, P=1.3×10^−3^) and being an athiest (OR=1.45, P=1.34×10^−2^) in EAs.

## Discussion

We employed both regression and non-regression based machine learning approaches to evaluate the association of more than 3,000 SUD, other psychiatric disorder, other health, and demographic variables with opioid cessation among EAs and AAs assessed in a cohort study of opioid, cocaine, or alcohol dependence. We observed moderate to high predictive accuracy across all methods, noting that SVM on average marginally outperformed the other methods. Although the specific set of predictive variables differed in EAs and AAs, a common profile emerged. People who ceased opioid tended to be older, initiated drug use later in life, had used opioids for a shorter period, experienced fewer problems related to cocaine or alcohol use, were currently employed, and had recovered from other psychiatric disorders including depression and PTSD.

Previous research using machine learning for addiction outcomes focused mainly on predictive accuracy, although a few studies attempted to identify and interpret specific variables that were associated with the outcomes(3, 4, 34, 35). Acion et al.(3) reported that ensemble super learning was superior to other machine learning methods, including penalized regression, SVM, and neural networks for predicting SUD treatment success indicated by treatment discharge status in a Hispanic cohort. In that study, less than 10% of participants had problems with cocaine or illicit opioids and fewer than 30 potential predictors were assessed. In contrast, we evaluated several thousand predictors, including detailed measures of drug-use activities and psychiatric disorders, and ranked the importance of the top-ranked variables with four distinct machine learning algorithms. Gowin et al.(3) identified regional brain activity changes predicting relapse from imaging data on fewer than 70 methamphetamine-dependent patients without including any lifestyle factors. Che et al.(4) applied deep learning to electronic health record data to predict people who used opioids short term, long-term, or with dependence. Similar to our study, they identified comorbid substance use and anxiety disorders as predictors(4). Several other studies used only regression-based methods to predict opioid and stimulant dependence(36), cocaine dependence(37) and alcohol dependence(38), which might not capture other relationships among variables. Several of the non-regression based methods we employed have also been applied in other studies, which focused mainly on MRI brain images as predictors of substance use disorder diagnoses(18, 19, 21, 34).

We identified several predictors of opioid cessation that were previously associated with other SUDs, including co-morbid drug use, antisocial behavior, suicidal thoughts, HIV infection, and asthma. Our finding that the majority of people who ceased opioids (60% in AAs and 66% in EAs) also ceased cocaine is consistent with evidence of high rates of co-occurrence of OUD and cocaine use disorder (CUD)(39, 40), which supports the development of treatment strategies to target both disorders (39, 41), and suggests that ceasing one substance might influence, or reflect, the ability to cease the other. Our findings are also consistent with observations that failure to address tobacco use lowers the efficacy of opioid cessation treatment(42) and that a behavioral intervention in patients with antisocial personality disorder reduces substance use(43). Unlike problems that are associated with other drug use, which predicts lower odds of opioid cessation, we found having cannabis use related problems predicts higher odds of opioid cessation (i.e had two marijuana symptoms lasting a month; marijuana interferes with work). This finding is puzzling and not immediately explainable. Previous findings of the co-occurrence of drug addiction, suicide attempts, depression, family conflicts, and PTSD, which may suggest bi-directional casual relationships,(44–49) are consistent with our observation that better management of comorbid psychiatric problems (fewer recent suicide attempts and psychiatric symptoms) increases the likelihood of opioid cessation.

One explanation for our findings of significant associations of ceasing opioids with a self-reported HIV diagnosis in AAs and asthma in EAs is that OUD patients with severe or life-threatening illnesses are more likely to seek or adhere to treatment,(50) an idea supported by evidence that HIV-infected OUD patients have better treatment outcomes for OUD.(51–53) Alternatively, poorer general health may lead to reduced drug use(54) (the so-called “sick quitter”).

We and Acion et al.(3) identified age, employment status, and age at first drug use as factors for treatment success. The protective effect of older age may be due to ascertainment bias because persons who survived severe dependence are more likely to have stopped using opioids. Further, our ascertainment strategy likely led to a higher proportion of older individuals and females who are in recovery from substance use disorders. Full-time employment likely reduces the time or urge for persons dependent on opioids to seek and use the drug. In addition, drug screening associated with some jobs may reduce the likelihood of current opioid use(55). The inverse correlation of age at first drug use and opioid cessation may reflect the increased difficulty to reverse the effect of long-term opioid exposure on the brain reward system(56), or increased severity associated with earlier onset. Other novel predictors of successful opioid cessation identified in this study among EAs include lower household atheism and income. The explanations underlying these associations are unclear, although a previous study showed that loss of religiosity between childhood and adulthood was associated with increased substance use while recent religiosity increased the odds of illicit drug use in the past year(57). In addition, lower household income might affect one’s ability to get opioids.

Implications can be drawn in terms of the overall accuracy from machine learning methods and individual factors identified. (A detailed discussion of the performance of the machine learning methods is provided in Supplemental Materials). Unlike standards established by a cardiologist reading an EKG or a radiologist reading a lung nodule, the accuracy as high as ~80% from our model has few comparable standards made by a physician specialized in substance misuse treatment. However, our accuracy result could serve as a reference for developing ways to address prescription opioid misuse and illicit opioid use, since we identified a solid list of individual variables to consider. While some of the factors identified were plausible and consistent with prior studies, factors like atheism and racing cars were less interpretable. Given that our research is atheoretical, results should be interpreted with caution and be validated.

The current study has several strengths. First, because the input dataset contains thousands of variables related to drug use activities, psychiatric disorders, medical history, and demographics obtained from several thousand individuals meeting DSM-5 criteria for mild to severe OUD, we were able to explore many factors in addition to those included in other studies. Second, both linear and non-linear machine learning methods were employed to best approach the true underlying model, which likely increased the robustness of our association findings. Third, we evaluated three models for each machine learning method to better understand the contribution of opioid and other drug use information. Finally, we considered only independent predictors in the association analyses to prevent over-representation of correlated factors.

Limitations of this work should be noted. First, given the cross-sectional nature of our data and the over 90% relapse rate(58) for OUD, many individuals classified as not using opioids may have subsequently relapsed to opioid use. However, it has been shown that prior abstinence is predictive of future abstinence.(59) Second, the machine learning analyses were based on samples that may have been underpowered to detect associations with some variables compared to other studies that included tens of thousands of individuals(60). Third, most persons in our cohort were evaluated prior to the current opioid epidemic and may not reflect recent secular trends in the prevalence and associated features of OUD. Fourth, associations of cessation with some variables and overall prediction accuracy may have been inflated because our analysis did not fully account for familial correlations. Fifth, in spite of the large number of variables that were included in the machine learning analyses, potentially important variables such as the reason for first use and details of treatment and support programs were unavailable. Finally, the rate of responses to many interview questions was substantially higher in EA participants, which may account for some of the observed racial differences in predictive models. Given these limitations, our findings require external validation in larger samples before they can be incorporated in prediction models for clinical purposes.

## Conclusions

Using machine learning techniques with feature selection, we analyzed a large number of variables include demographic, behavioral, health and drug use activities and found variables in a wide range of domains that were associated with cessation. These included some that are consistent with prior literature, others that plausibly could be associated but have not been well-studied, and others that do not have readily apparent explanations for their associations. These findings suggest hypotheses for future studies and could inform how one might increase the likelihood of cessation with and without treatment. The findings support a number of widely known treatment strategies for OUD, such as treating psychiatric comorbidity, adding wraparound services like employment counseling, and simultaneously addressing polydrug use problems. Finally, in an era of increasing availability of digitized health-related records, our study provides a framework for disease outcome prediction using high dimensional health-related records.

