## Supplemental figures for "Factors associated with opioid cessation: a machine learning approach"

### Supplemental Methods

Least Absolute Shrinkage and Selection Operator (LASSO)

LASSO implemented in the R package ‘glmnet’(1) was used for both feature selection and prediction. The shrinkage parameter lambda in the penalty term of LASSO regression was obtained using 10-fold cross validation on the training set 10 times. Separate accuracy criteria of either misclassification error or AUC were used to search for the lambda with the best model fit. The “1SE rule(2)” which aims to find the simplest model with comparable accuracy to the best model, was used to identify lambda that associated with the lowest cross validation error as well as lambda whose cross validation error was one standard error unit from the lowest cross validation error on the training set. We identified the significant features by fitting lambdas on the training set. The test set accuracy was evaluated by using the class probability prediction of test set as an input to Scikit-learn(3) (roc_auc_score and f1_score ) to obtain our final accuracy measures: F1 score and AUC.

Support Vector Machine (SVM) with Recursive Feature Elimination

Linear SVM with recursive feature elimination was implemented using the Scikit-learn python package. A soft margin was included in the model to reduce overfitting. Briefly, we set the penalty parameter C by exhaustively searching the range of values between $2^{-20}$ and $2^{8}$ that covers the recommended range proposed by Hsu et.al(4) using 10-fold cross validation with balanced class weights on the training set . We applied the parameter C chosen by either maximizing AUC or F1 score (whichever yielded the highest accuracy) in the training set for recursive feature selection. A feature’s importance, represented by its weight, was used as the input to the recursive feature elimination function, where 10-fold cross validation was used to find the combination of features that maximizes either AUC or F1 score (whichever yielded the highest accuracy). Selected features from each model were applied to the test set to obtain the overall test accuracy.

Random Forest (RF) Recursive Feature Elimination

The RF recursive feature elimination approach was implemented using the Scikit-learn python package. We determined the ideal number of decision trees needed for the RF model by searching the recommended range of $2^7$ to $2^{11}$ proposed by Oshiro et.al(5) 10-fold cross validation with balanced class weight on the training set. The decision trees that associated with the maximum AUC or maximum F1 score were individually chosen. A feature's importance, represented by the Gini impurity(6) measure proposed by Breiman(7), was used to evaluate the importance of a variable by adding up the weighted impurity decreases for all nodes averaged over all decision trees. A feature's importance score??? was used as the input for recursive feature elimination and 10-fold cross validation was used to find the combination of features that maximized either AUC or F1 score (whichever yielded the highest accuracy). Selected features from each model were included in the test set without any feature selection steps to obtain the test accuracy.

Deep Neural Network (DNN)

The DNN approach was applied using Keras(8) with a tensorflow framework. A 3-layer, fully connected feed forward DNN was constructed with two hidden layers using rectified linear units (Relu) as an activation function and a sigmoid function for the output layer. The training set that was used in the aforementioned methods was further split into a development set and validation set with 9:1 ratio with fixed case control ratios to minimize overfitting or inadequate training. The Adam optimization method was used to find efficiently the parameters associated with the ideal state of the DNN objective function. The Adam optimizer(9) has been proven to be the start of art(10-12) optimization method by incorporating the advantages of two most popular optimizations (RMSProp(13) and AdaGrad(14)). We followed the suggested hyperparameter settings from the developer of the Adam optimizer(9) and only tuned the learning rate. L2 normalization with various scales were performed on each layer to prevent overfitting. Cross validation of the development set and balanced accuracy were used to reduce bias for the hyperparmeter search. Both weighted AUC and weighted F1 score on the validation set were used to measure model performance.

Several methods have been proposed for feature selection using DNN, with the focus on reducing input dimensionality, such as sparse one-to-one, dropout feature ranking, and activation potential based(15-17). We used the activation potential based method because of its proven performance in reducing the number of features, to obviate application of another filtering method coupled with DNN, and its simplicity and intuitiveness for selecting the number of important variables. Feature selection was performed according to the method proposed by Roy et.al(17). Briefly, we computed the activation potential of each input feature connected to each of the hidden nodes in the first layer before applying Relu. Then, the average relative activation potential of each feature in each first hidden layer node was calculated by averaging the number of input features and training samples at each node. Relu activation was applied to the average relative activation potential to obtain the net positive contribution of each input feature. Input features were ranked and plotted against their net positive activation potential contribution. Important features were chosen based on their collective contribution of net positive activation potential. Because of bias associated with AUC in the presence of imbalanced dataset, the feature combination that associated with the highest F1 score in the validation set was used to obtain the accuracy on test set.

Performance of Machine Learning Algorithms

There was a consistent drop in accuracy from model 1 to model 3 across the four machine learning methods, although the difference between models 1 and 2 is smaller than the difference between models 2 and 3 (Figure S2). The loss of accuracy across models was greater in AAs than in EAs.  Thw AUC generally demonstrated higher accuracy than the F1 score. Among the machine learning methods, SVM yielded the highest F1 score more frequently than the other methods across models in both AAs and EAs. Most notably, SVM had the best performance for model 2 in AAs and models 2 and 3 in EAs, although the differences in accuracy between SVM and the other models with high performance were small. The observation that SVM performed only marginally better than LASSO in both AAs and EAs was not surprising because SVM using a linear kernel and LASSO employ a linear model with regularization. Both SVM and LASSO selected uncorrelated features, however SVM also selected correlated features. Both RF and DNN showed some evidence for overfitting (result not shown), although the effect was relatively small that was reflected by an approximately 4% higher accuracy in the cross validation training set than the test set. The RF model may have been overfitted because the number of individual classifier decision trees was fixed. DNN generally requires a much larger sample size than the one available here, which might have limited its performance.

**Supplemental Table 1**. Five most significant variables ranked by feature importance for each machine learning method, stratified by model and population.

African Americans

| | | Variable | Rank | p-value |
|---|---|---|---|---|
| Model 1 | LASSO | Recency of last cocaine use | 1 | 2.89E-07 |
| | | Current age | 2 | 5.39E-10 |
| | | Recency of 1st opioid treatment | 3 | 4.97E-06 |
| | | Cocaine use severity | 4 | 4.69E-03 |
| | | N years using Heroin | 5 | 2.80E-08 |
| | SVM | Recency of last cocaine use | 1 | 3.50E-08 |
| | | Recency of last cocaine injection | 2 | 1.38E-04 |
| | | Current age | 3 | 9.20E-07 |
| | | N years using Heroin | 4 | 7.68E-06 |
| | | Recency of weekly cocaine use for 1 month | 5 | - |
| | RF | Recency of last cocaine use | 1 | 4.38E-06 |
| | | Recency of weekly cocaine use for 1 month | 2 | - |
| | | Used cocaine>11 last year | 3 | - |
| | | Didn't use cocaine>11 last year | 4 | - |
| | | Recency of cocaine symptoms | 5 | - |
| | DNN | Recency of last cocaine use | 1 | 1.04E-06 |
| | | Recency of last cocaine injection | 2 | - |
| | | Recency of cocaine symptoms | 3 | 1.11E-01 |
| | | Current age | 4 | 5.50E-08 |
| | | Recency of weekly cocaine use for 1 month | 5 | - |
| Model 2 | LASSO | Recency of last cocaine use | 1 | 5.32E-15 |
| | | Current age | 2 | 3.96E-08 |
| | | Cocaine use severity | 3 | 1.24E-07 |
| | | Recency of last tobacco use | 4 | 1.62E-03 |
| | | Recency first reported drug problems to professionals | 5 | 9.65E-01 |
| | SVM | Recency of last cocaine use | 1 | 5.44E-07 |
| | | Recency of weekly cocaine use for 1 month | 2 | - |
| | | Recency of cocaine symptoms | 3 | - |
| | | Didn't use cocaine>11 last year | 4 | 2.61E-02 |
| | | Used cocaine>11 last year | 5 | 5.16E-02 |
| | RF | Recency of last cocaine use | 1 | 2.26E-06 |
| | | Used cocaine>11 last year | 2 | - |
| | | Didn't use cocaine>11 last year | 3 | 4.38E-02 |
| | | Recency of cocaine symptoms | 4 | - |
| | | Recency of weekly cocaine use for 1 month | 5 | - |
| | DNN | Age first used tobacco | 1 | 9.42E-04 |
| | | Had relationship for >1 year | 2 | 6.54E-03 |
| | | Longest time in days without using tobacco | 3 | - |
| | | Contacted relatives infrequently | 4 | - |

| | | | | |
|---|---|---|---|---|
| | | Mixed drugs with alcohol >3 times | 5 | - |
| Model 3 | LASSO | Current age | 1 | 5.20E-11 |
| | | HIV positive | 2 | 8.55E-06 |
| | | Blamed others for one's mistake | 3 | 1.76E-03 |
| | | Jobless for 6 month due to drugs/alcohol | 4 | 2.93E-02 |
| | | Currently unemployed | 5 | 2.97E-04 |
| | SVM | Current age | 1 | 2.67E-09 |
| | | N biological children | 2 | 3.62E-02 |
| | | Currently unemployed | 3 | 3.94E-02 |
| | | Age at maximum weight | 4 | - |
| | | N months employed last year | 5 | - |
| | RF | Current age | 1 | 2.49E-09 |
| | | Age at maximum weight | 2 | - |
| | | BMI | 3 | 5.98E-06 |
| | | N drug symptoms when having depression | 4 | 1.13E-02 |
| | | current weight | 5 | - |
| | DNN | N doctor visits for health problems | 1 | - |
| | | Current age | 2 | 5.17E-11 |
| | | Jobless for 6 month due to drugs/alcohol | 3 | 1.49E-01 |
| | | Treated for emotional, psychiatric or drug problems | 4 | 3.51E-03 |
| | | Visited treatment center once last year | 5 | 1.25E-01 |

European Americans

| | | Variable | Rank | p-value |
|---|---|---|---|---|
| Model 1 | LASSO | Recency of last cocaine use | 1 | 8.72E-19 |
| | | Current age | 2 | 9.96E-12 |
| | | Recency started opioid treatment | 3 | 9.31E-07 |
| | | Recency of last cocaine injection | 4 | 1.80E-04 |
| | | Age last had antisocial behaviors | 5 | 3.92E-06 |
| | SVM | Recency of last cocaine use | 1 | 1.01E-16 |
| | | Recency of last cocaine injection | 2 | 1.01E-06 |
| | | Used cocaine>11 times in last year | 3 | 3.10E-03 |
| | | Age at maximum weight | 4 | 5.46E-02 |
| | | Current age | 5 | 9.87E-03 |
| | RF | Recency of last cocaine use | 1 | 4.38E-06 |
| | | Recency of cocaine symptoms | 2 | 3.24E-03 |
| | | Didn't used cocaine>11 times in last year | 3 | 2.79E-03 |
| | | Recency of last cocaine injection | 4 | 6.52E-03 |
| | | Used cocaine>11 last year | 5 | - |
| | DNN | Recency of last cocaine use | 1 | 4.79E-25 |
| | | Recency first brought up opioid problems with professionals | 2 | - |
| | | Recency of last cocaine injection | 3 | 1.07E-03 |
| | | Age last smoked cigarettes | 4 | 6.60E-21 |
| | | No drunk driving arrests | 5 | - |
| Model 2 | LASSO | Recency of last cocaine use | 1 | 3.06E-18 |

| | | | | |
|---|---|---|---|---|
| | | Current age | 2 | 1.27E-05 |
| | | Recency of last cocaine injection | 3 | 1.38E-08 |
| | | Age last had antisocial behaviors | 4 | 4.56E-06 |
| | | Cocaine use severity | 5 | 1.83E-07 |
| | SVM | Recency of last cocaine use | 1 | 3.14E-17 |
| | | Recency of last cocaine injection | 2 | 1.74E-06 |
| | | Didn't use cocaine>11 last year | 3 | 8.36E-05 |
| | | Age at maximum weight | 4 | 3.82E-02 |
| | | Current age | 5 | 1.64E-05 |
| | RF | Recency of last cocaine use | 1 | 3.13E-14 |
| | | Recency of last cocaine injection | 2 | 2.43E-06 |
| | | Used cocaine>11 last year | 3 | - |
| | | Didn't use cocaine>11 last year | 4 | 8.16E-04 |
| | | Recency of cocaine symptoms | 5 | - |
| | DNN | Recency of last cocaine use | 1 | 1.42E-17 |
| | | Didn't hurt animal on purpose | 2 | - |
| | | Recency last stayed high from cocaine >1 day | 3 | 2.83E-03 |
| | | Recency had >2 cocaine symptoms | 4 | - |
| | | Recency had cocaine symptoms | 5 | - |
| Model 3 | LASSO | Current age | 1 | 2.36E-09 |
| | | Current health has always been worse | 2 | 7.62E-07 |
| | | BMI | 3 | 3.12E-13 |
| | | Age last had antisocial behaviors | 4 | - |
| | | Depression started with drug problems | 5 | 3.98E-03 |
| | SVM | Current age | 1 | 3.10E-08 |
| | | Age at maximum weight | 2 | 4.62E-03 |
| | | Age last had antisocial behaviors | 3 | 2.74E-07 |
| | | BMI | 4 | 5.89E-12 |
| | | Current weight | 5 | - |
| | RF | Current age | 1 | 5.10E-09 |
| | | Age at maximum weight | 2 | 2.36E-02 |
| | | BMI | 3 | 1.15E-04 |
| | | Current weight | 4 | 8.08E-02 |
| | | Max weight | 5 | 4.51E-03 |
| | DNN | Current age | 1 | 2.13E-09 |
| | | Bad mood after ECT or bright light therapy | 2 | - |
| | | Didn't have OCD | 3 | - |
| | | No gambling withdrawal when cannot gamble | 4 | 9.20E-02 |
| | | Didn't have sex with >10 people in a year | 5 | 3.54 E-02 |

"Rank": relative importance of a variable measured by the specified method; "-" : variable not selected by stepwise regression; LASSO: least absolute shrinkage and selection operator; SVM: support vector machine with recursive feature elimination; RF: random forest with recursive feature elimination; DNN: deep neural network with recursive feature elimination.

**Supplemental Figure 1**. Derivation of African Americans (AAs) and European Americans (EAs) subjects in the Yale-Penn dataset who were ascertained from multiple substance use disorder studies and met criteria for cessation or non-cessation of opioid use.

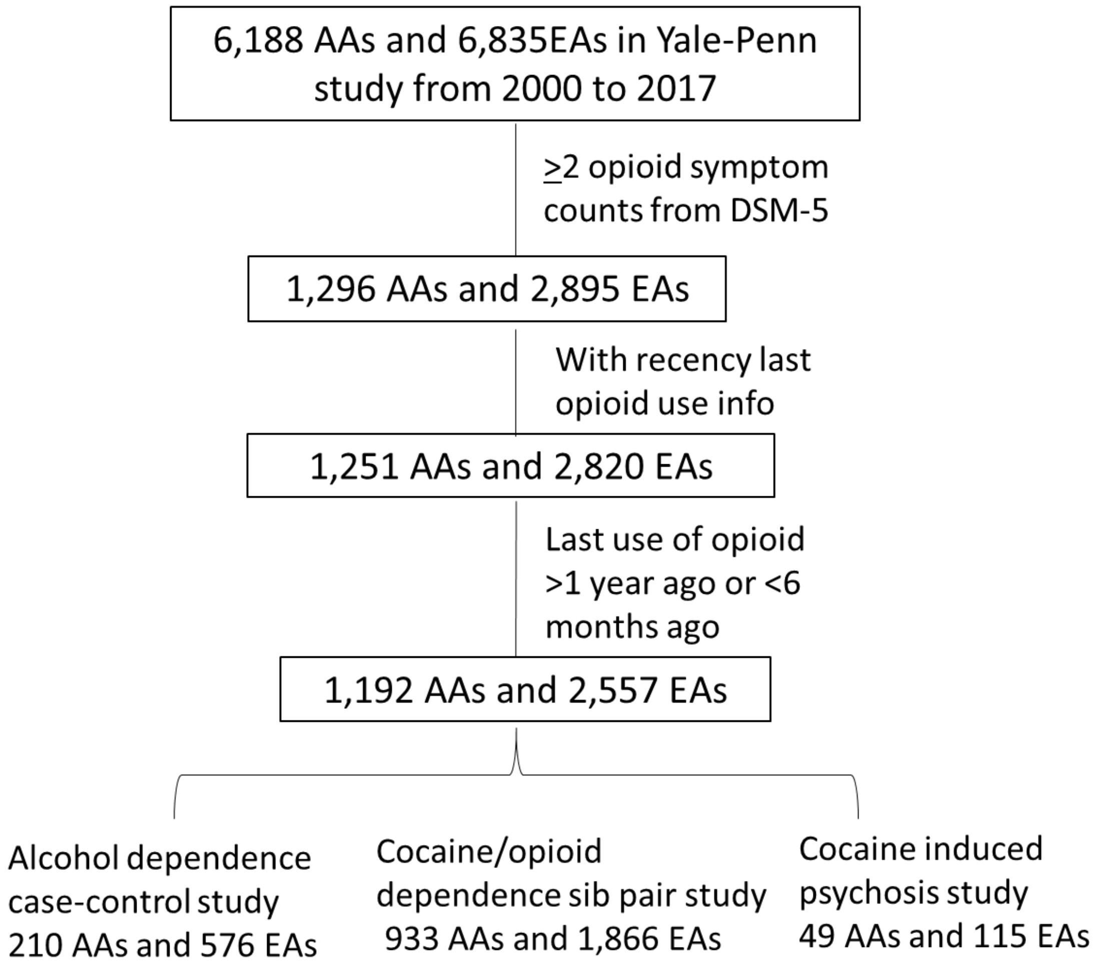

**Supplemental Figure 2**. Predictive accuracy of four machine learning methods by model in African Americans (A) and European Americans (B). AUC and F1 scores derived from a common set of variables picked by the method (DNN, LASSO, RF, SVM) for each model. m1: model 1; m2: model 2; m3: model 3; DNN: deep neural network; LASSO: least absolute shrinkage and selection operator; RF: random forest; SVM: support vector machine; AUC: area under receiver operating curve; F1: F1 score.

A)

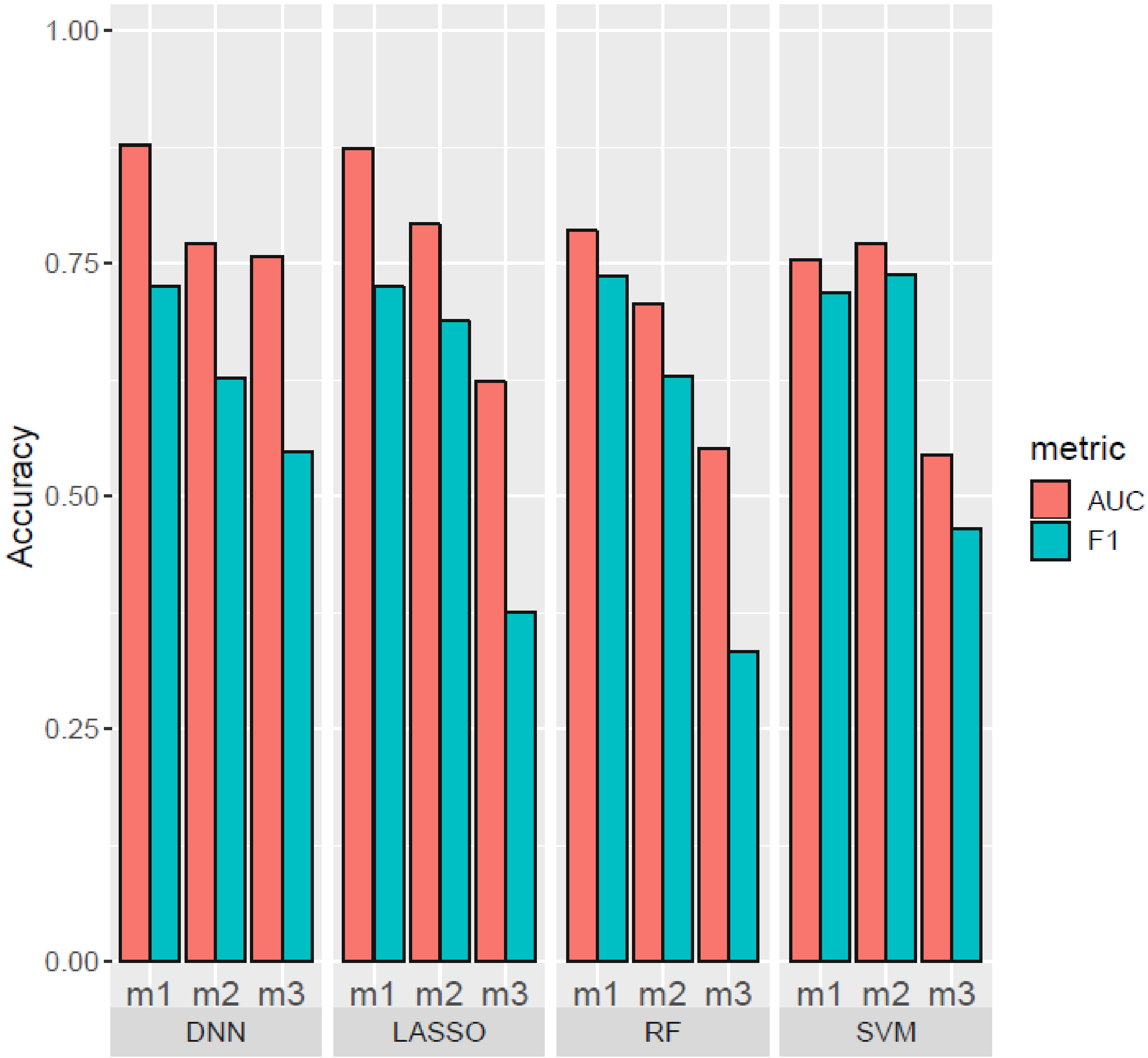

B)

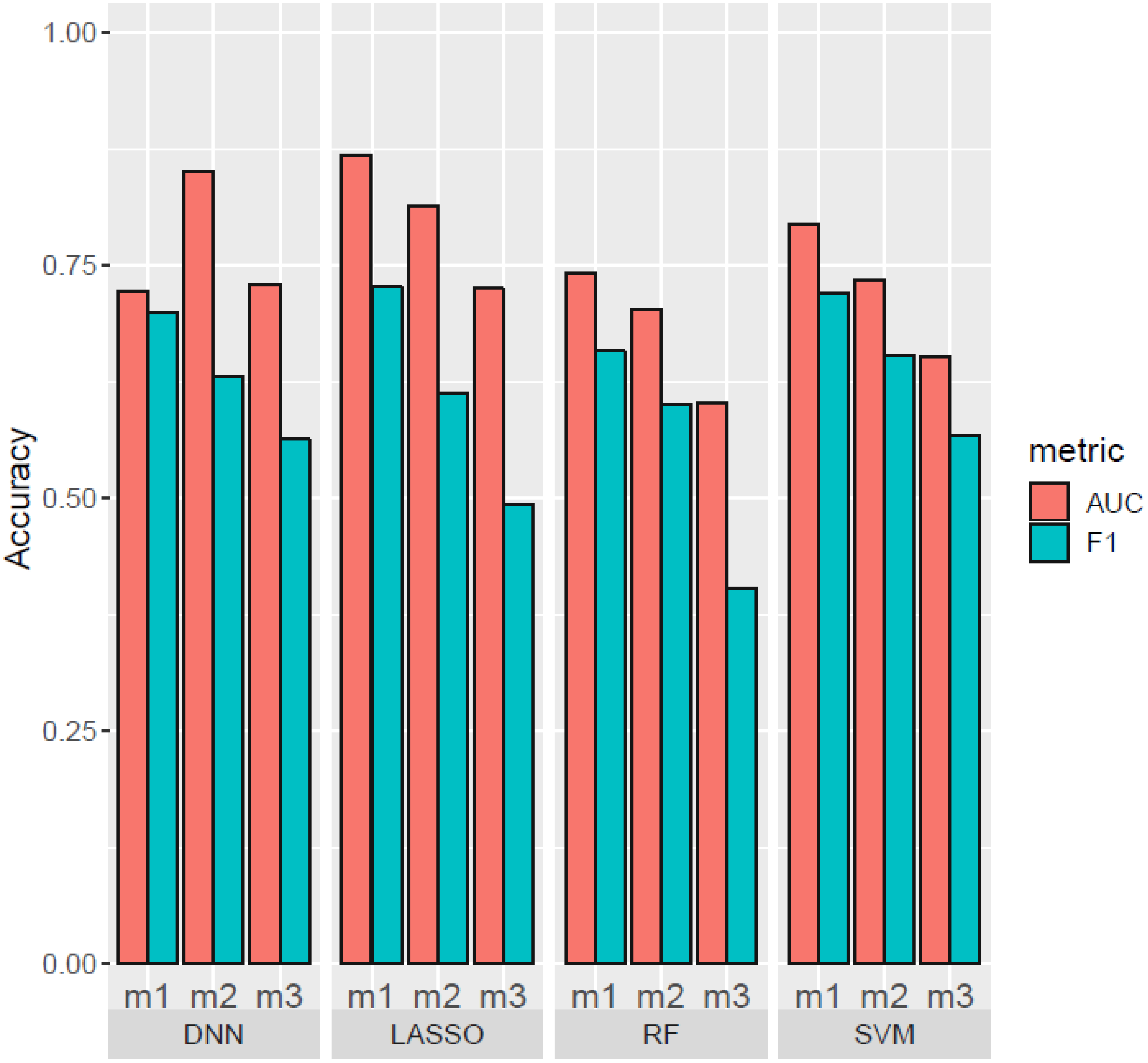

1.00
0.75
0.50
0.25
0.00
Accuracy
m1
m2
m3
DNN
LASSO
RF
SVM
metric
AUC
F1

**Supplemental Figure 3.** Number of overlapping “high impact” variables selected by each machine learning method based on the importance measurement in African Americans (A) and European Americans (B) for models (1), (2), and (3). Model 1 includes all variables except for those that are confounded with opioid cessation. Model 2 includes all variables in Model 1 except opioid-related variables. Model 3 includes all variables in Model 2 except drug related variables. The criteria used for ‘high impact’ variables of each machine learning method can be found from **Materials and Methods** section in the main text. Colors represent machine learning method: pink = LASSO, light yellow = SVM, light green =RF, light blue = DNN.

A)

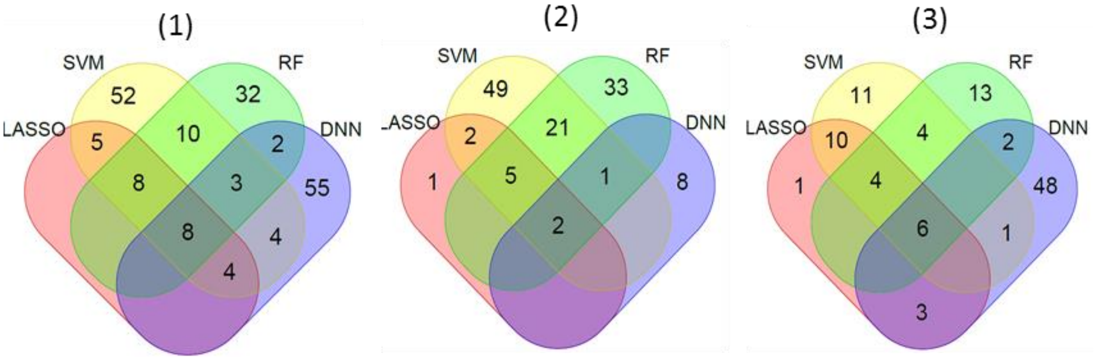


B)

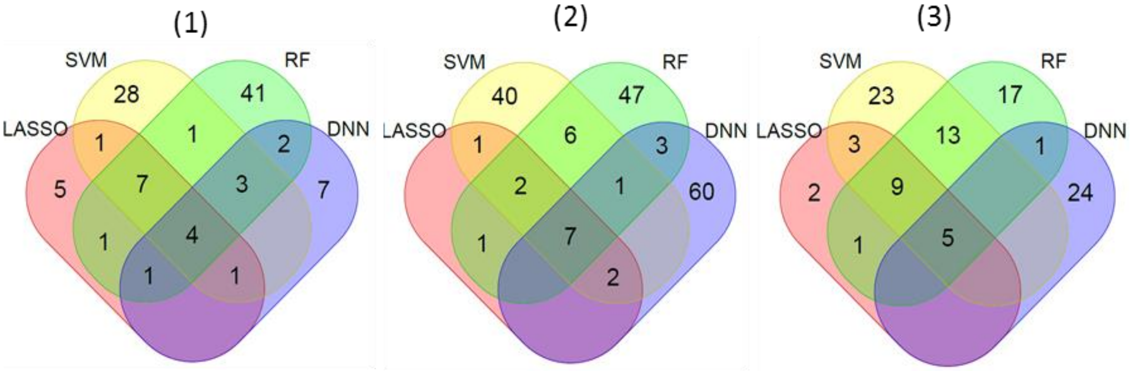
